# Entering into a self-regulated learning mode prevents detrimental effect of feedback removal on memory

**DOI:** 10.1101/2021.07.02.450865

**Authors:** Peter Vavra, Leo Sokolovič, Emanuele Porcu, Pablo Ripollés, Antoni Rodriguez-Fornells, Toemme Noesselt

## Abstract

Recently, we provided causal evidence that self-regulated dopamine signaling enhanced long-term memory formation in the absence of any external feedback or reward (Ripollés et al., 2016, 2018) if a congruent meaning inferred from semantic context (DA-dependent learning), while DA-signals were absent if no congruent meaning could be inferred (DA-independent learning). Here, we investigated the interaction of self-regulated learning with externally-regulated DA-signalling by providing external performance feedback in the first or second half of trials. We found that removing feedback during DA-dependent learning lowered subsequent recognition rates a day later, whereas recognition remained high in the group which received feedback only in the second half. In contrast, feedback modestly enhanced recognition rates for both groups for DA-independent learning. Our findings suggest that external reinforcers can selectively impair DA-dependent memories if internal DA-dependent processes are not already established and highlights the relevance of self-regulated learning in education to support stable memory formation.

## Introduction

The interplay of internal motivational states and external incentives governs our behavior (Bénabou & Tirole, 2003; Dweck, 1986; Ryan and Deci, 2000; Camerer & Hogarth, 1999). Usually, intrinsic motivation refers to those activities that are engaged for their very own sake (e.g., enjoyment and pleasure) and are not necessarily associated with a particular instrumental and external benefit (e.g., incentives). (for review see e.g. Cerasoli et al., 2014). A key finding, the *undermining effect* (for a review, see Deci, Koestner, Ryan, 1999), posits that the introduction of extrinsic incentives “crowds out” intrinsic motivation. Moreover, performance plummets when the incentives are subsequently removed. While there has been some controversy about the robustness of this effect (e.g. Deci, Koestner, Ryan, 2001; Cameron, 2001), a recent quantitative meta-analysis (Cerasoli et al., 2014) found evidence for this effect: explicit rewards increase task performance, but reduce the influence of intrinsic motivation on the task outcome. A determining factor accounting for previous differences across studies appears to be the use of distinct performance measures (e.g., qualitative vs quantitative). In educational contexts, however, there is another crucial dimension to consider: do such motivational factors affect only online performance (e.g., how well one understands the meaning of a new-word when first encountering it), or does the interplay between different motivational factors also influence longer-lasting memory processes (e.g., remembering the new vocabulary and reducing natural forgetting)?

Neurobiologically, there is accumulating evidence showing that external rewards can boost learning and memory performance by activating a dopaminergic loop mainly formed by the substantia nigra/ventral tegmental area complex (SN/VTA), the hippocampus (HP) and the ventral striatum (VS; hereafter referred to as SN/VTA-HP loop; Lisman and Grace, 2005; Goto and Grace, 2005; Lisman et al., 2011; Shohamy and Adcock, 2010). The loop is activated when new information arrives at the HP, which sends a signal to the VS where affective, motivational, and goal-directed information is integrated (Lisman and Grace, 2005; Goto and Grace, 2005). The VS can then disinhibit dopamine-releasing neurons in the SN/VTA complex which project back to the HP. The release of dopamine back at the HP induces Long-Term Potentiation processes (LTP) which ultimately enhance long-term memory storage.

In addition to external primary and secondary reinforcers like food or money, external performance feedback can also boost activity in SN/VTA and VS (e.g. Ullsperger & Cramon, 2003), as well as long-term memory (Fazio and Marsch, 2009). Importantly for our work, this SN/VTA-HP loop can also be modulated in absence of any external signal, e.g. by intrinsic states like curiosity (Gruber et al., 2014; see review Gruber & Ranganath, 2019), or by self-regulated learning (Ripollés et al., 2014; 2016). In our previous work, participants learned the meaning of new-words from a semantic context formed by two sentences (Ripollés et al., 2014; 2016; 2018). The meaning in the two sentences were congruent for half of the new-words and, thus, the second sentence disambiguated between different meanings (e.g. *“Every Sunday the grandmothers went to the jedin”* and *“The man was buried at the jedin”* (likely congruent meaning: graveyard)); for the other half of the new-words, the sentences were incongruent (i.e., no coherent meaning could be derived, e.g*. “Every night the astronomer watched the heutil”* (likely meaning: moon, stars or some other extraterrestrial object) and “*Every morning the coworkers drink heutil*” (likely meaning: coffee, tea)). Participants were told to extract the meaning of the new-word when the meaning was congruent and to detect the incongruence otherwise. Critically, we showed that, in absence of any explicit reward, successful learning was tightly coupled with increased self-reports of pleasure, enhanced electrodermal activity and, most importantly, heightened fMRI activity in the SN/VTA-HP loop (Ripollés et al., 2014, 2016). Moreover, behaviourally intrinsically generated reward-related signals paralleled an increase in memory retention for these new-words in a surprise memory test assessing long-term memory on the next day and was still detected when testing recognition memory after 7 days (Ripollés et al., 2016). The successful discovery of the incongruence between the meanings evoked by the two sentences was neither related to a heightened behavioral or physiological measure, nor to increased reward-related activity in the SN/VTA-HP loop. In our subsequent work (Ripollés, et al., 2018), we employed a double-blind, within-subject randomized pharmacological manipulation of the dopaminergic system. Our results confirmed that dopaminergic signaling had a causal role in improving performance during the initial learning phase, as well as long-term memory for the new-words learned from congruent contexts (i.e. DA-dependent motivated-learning) while we did not observe a significant relationship for incongruent contexts (i.e. DA-independent learning).

Given that both extrinsic feedback and intrinsic learning signals appear to be DA-dependent and apparently activate at least parts of the same reward-related SN/VTA-HP loop, a crucial question is how these two processes interact or to which extent they compete (Murayama et al., 2014; Deci et al., 1999; Morgan, 1984; Lepper et al., 1973). Here, we extend our prior research by adding trial-based feedback on participants’ accuracy in our new-word learning paradigm (Ripollés et al., 2014, 2016, 2018). Performance feedback externally triggers the SN/VTA-HP loop in a similar fashion as primary rewards and could provide a more ecologically valid setting for learning paradigms in humans than presenting primary or secondary reinforcers. Our aim was to investigate how two distinct DA-dependent processes – external feedback and intrinsic self-regulated learning – interact. To this end and based on our previous work, we compared learning trials involving intrinsically triggered dopaminergic activations (discovering a meaning from a congruent context, DA-dependent) with trials where such an intrinsically triggered DA-signal is absent (incongruent contexts, DA-independent). Crucially, the inclusion of both congruent and incongruent trials allows us to study the effect of feedback on two different learning processes using well-matched stimuli within the same paradigm, where one process relies on the dopaminergic SN/VTA-HP loop and therefore on DA transmission, and the other is operating independently of it. We hypothesized that task performance should be affected by adding external feedback, but that this effect would differ depending on the two trial-based internal DA-milieus in our experiment. During congruent trials, an interplay of DA-dependent external feedback with DA-dependent intrinsic signals should be observed since dopaminergic signaling is directly related to learning performance and memory consolidation for this condition (Ripollés et al., 2018). In contrast, during incongruent trials the task is to learn that the meanings between the two sentences were incongruent and to store this more general association in memory (i.e., that a particular new-word has an incongruent meaning). Further, we hypothesized that the timing is critical at which external feedback is introduced. According to the undermining effect, feedback which was presented from the beginning of the task should decrease performance once it is removed. Althought not implied by the undermining effect, adding feedback after an initial feedback-free phase might increase also performance. Finally, we assessed whether feedback and its timing affects performance during learning and memory differently.

## Results

Sixty healthy participants completed an altered version of our word-learning task (see also Materials and Methods and Fig. 1). In this task, participants read sentences which end in a new-word. Importantly, each new-word was presented at the end of two sentences and these pairs of sentences could be congruent (M+ trials; e.g. “Every Sunday the grandmother went to the *jedin*” (sentence 1); “The man was buried at the *jedin*” (sentence 2); congruent meaning: graveyard), allowing the inference of a coherent meaning of the new-word; or sentences could be incongruent (M-trials; e.g. “I bought tickets for the *jardy*”; “The workers drink coffee with *jardy*”; meaning 1: concert/football game etc. meaning 2: milk, sugar, alcohol), such that no unique meaning fits both sentences. After reading the second sentence and the associated new-word, participants verbally responded by either saying what the congruent meaning of the new-word had been, or indicating that the sentences had been incongruent. Finally, participants could also indicate that they were unsure by remaining silent or by saying “‘I don’t know’”. In contrast to our previous work, in the current version of the task we provided participants with trial-based informative performance feedback on half the trials (smileys, frownies) or showed them non-informative placeholders pictures on the other half (scrambled smileys/frownies; here-after called “no feedback”). After approximately 24 hr, participants completed a surprise memory test to assess their learning: Thirty-one participants completed a free-recall memory test where they had to indicate which new-words they remembered and indicate their meaning, or that the two sentences had been incongruent. The other twenty-nine participants completed a recognition test (chance level was 25%; see Materials and Methods for details). During the learning and recognition session participants also rated pleasantness, arousal and confidence after each trial, in order to assess the subjective states underlying the behavioral responses (s. Fig. 1).

**Figure 1:**
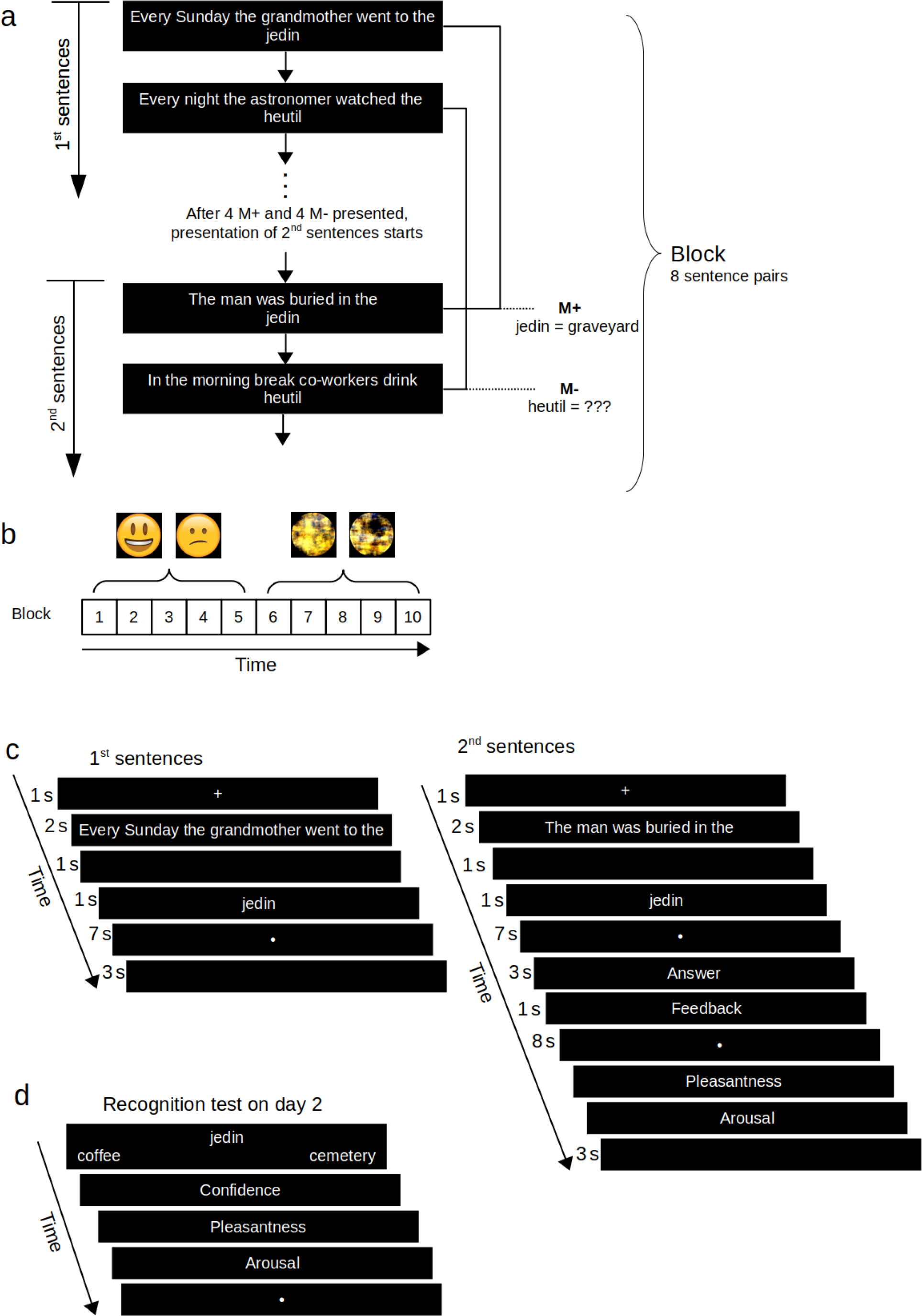
Experimental Procedure. **A.** Example of a block with 8 first and 8 second sentences. In each block 4 M+ and 4 M-pairs of sentences were presented. In the first half of each block all eight first sentences were presented, all ending in a different new-word (see above for one M+ and M-example sentence pair each). After the presentation of all 8 first sentences, the second sentences for those same new-words were shown in a randomized order relative to the first sentences. **B.** Exemplary overall block structure of the experiment for one participant who started with feedback (blocks 1-5) and switched to no feedback in the second half (block 6-10). Across participants feedback order was counterbalanced, i.e. feedback was present in the first or second half of the blocks (feedback-1.half or feedback-2.half). All participants received feedback in form of a smiley or frownie and scrambled smileys/frownies (no feedback). **C.** An exemplary M+ trial in the learning session adapted from Ripollés et al. (2016). In each paired learning trial, two sentences ending in the same new-word were presented. For M+ trials, the two sentences evoke the same congruent meaning (“graveyard”) and thus DA-dependent word-learning can occur (i.e., *jedin* means “graveyard”). M-trials follow the same structure with the difference that the two sentences do not evoke the same meaning (i.e., participants can only learn that this new-word had an two incongruent meanings, DA-independent). After the presentation of the new word, subject’s response and feedback, subjective ratings of pleasantness and arousal were obtained. **D.** An example of a recognition test trial on day 2. All new-words (both M+ and M-) from the learning session were presented in a randomized order with two possible meanings to choose from. In addition, participants could indicate that the meaning for that new-word had been incongruent, or that they didn’t know. After indicating their answer, participants again rated their pleasantness and arousal, as well as confidence.

### Adding external feedback improves immediate performance on day 1

We first assessed whether external feedback influenced immediate task performance (Figure 2a) on day 1 using a generalized linear mixed model with the within-subject factors feedback (present/absent) and congruence (M+/M-) and the between subject-factor order (feedback first/later).

**Figure 2:**
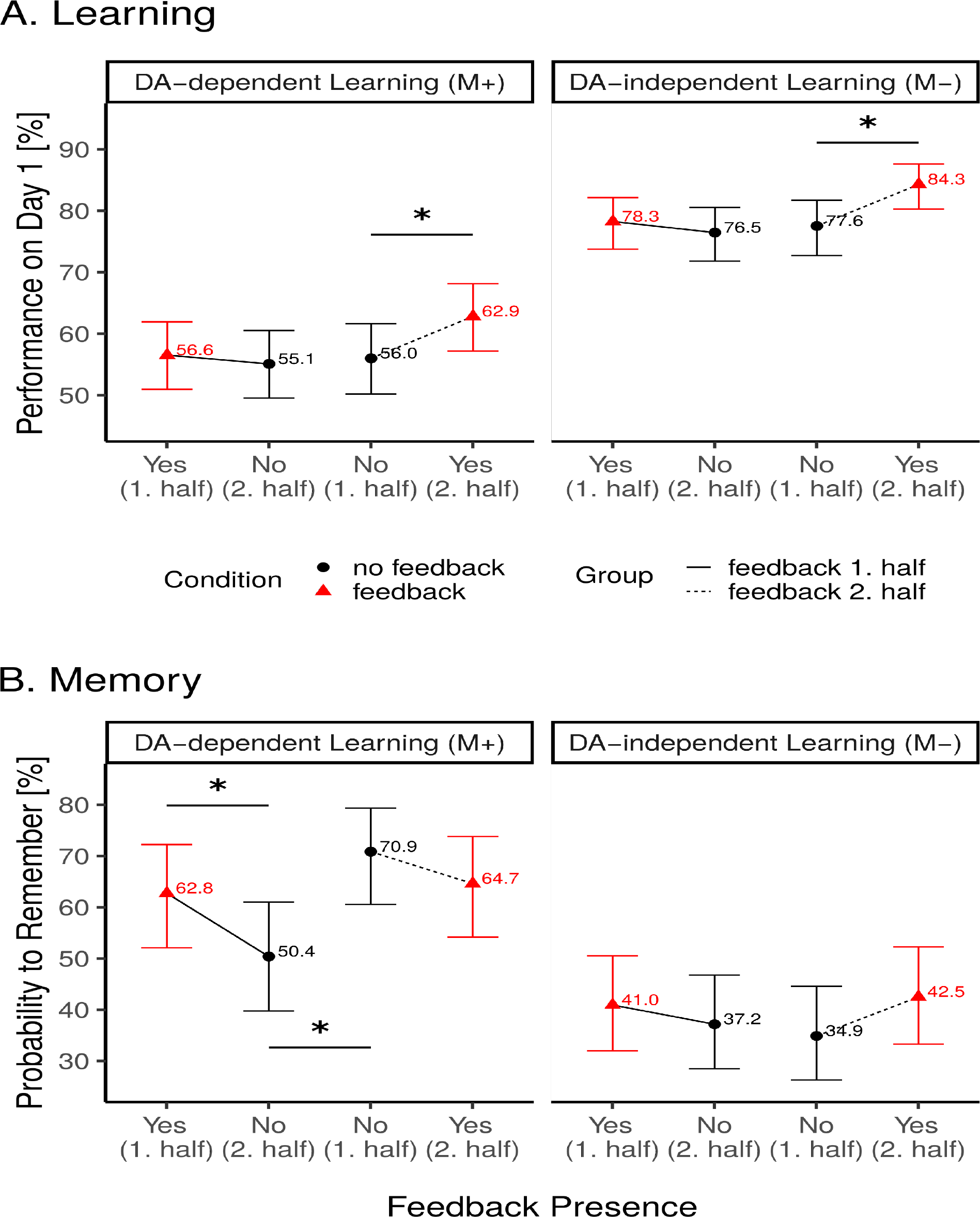
Performance on new-word meaning extraction task on day 1 (A) and recognition memory on day 2 (B). Trials with no congruent meaning (M-) are displayed on the right, trials with congruent meaning (M+) on the left, separately for feedback order (between subject effect; lines connect data points for the feedback-first half and feedback-second half groups; full line: started with feedback; dotted line: started without feedback) and feedback presence (within subject effect; red triangle: feedback; black dot: no feedback). Note that feedback presence is plotted in order of occurrence on x-axis (i.e. reversed feedback presence for the two groups). Lines between data points indicate identical within-group measures differing only in feedback presence. Error bars indicate 95% confidence interval. **A. Immediate performance during meaning extraction task (implicit learning).** Across conditions M-trials (right side) showed a higher level of performance than M+ trials. Informative feedback improved performance for the group of participants who started off without feedback and started receiving feedback on their answers in the second half of the experiment. Importantly, this pattern was observed for both M+ and M-trials (p < .023 and p < .004 for M+ and M- trials respectively). **B. Recognition Performance on day 2** as a function of congruence of meaning across sentences, feedback order and feedback presence (chance-level = 25%). In contrast to day 1, incongruent new-words (M-) were less likely to be remembered than congruent new-words (M+). For incongruent trials, there was a trend for feedback boosting recollection rates. For congruent trials, removing feedback in the second half of the experiment lead to a substantial drop in recollection rates a day later (p< .034), whereas no effect was found if feedback was added in the second half of trials (i.e., if participants started the learning phase with no-feedback, p< .24).

On average, participants were able to correctly infer the meaning of new-words from the congruent contextual information (congruent trials: 57.7±1.7% correct, see Figure 2; this learning rate is in accord with our previous work, Ripollés et al., 2014, 2016, 2018). They were also able to correctly indicate when two sentences were incongruent (incongruent trials: 79.4±1.4% correct; main effect of congruence: χ^2^(1) = 248.6, *p* < .0001, see Fig. 2a). Importantly, receiving feedback improved the overall performance during the implicit learning phase (4.7% difference; main effect of feedback: χ^2^(1) = 10.9, *p* = .001). However, this effect was mostly driven by the order in which feedback was provided: When participants who began without feedback started receiving feedback in the second half of the experiment, their performance improved substantially (67.7% to 75.1% when collapsing across incongruent and congruent trials, odds-ratio=0.696±0.069, *p* = .0003); the corresponding drop in performance for the group of participants for whom the feedback was withdrawn was substantially smaller (68.4% with feedback to 66.7% without, odds-ratio=0.923±0.083, *p* = .37; interaction feedback-by-order: χ^2^(1) = 4.42, *p* = .04). These effects of feedback were not significantly different for the congruent/incongruent trials (congruence-by-feedback: χ^2^(1) = 0.57, *p* = .45, congruence-by-order: χ^2^(1) = 0.36, *p* = .55; congruence-by-feedback-by-order: χ^2^(1) = 0.18, *p* = .67).

Overall, present results showed that introducing feedback in the second half of the experiment improves online word learning performance, whereas the withdrawal of feedback led to no significant drop in online-performance on day 1 (see Fig. 2a).

### Removing external feedback decreases DA-dependent long-term memory

To assess how this external feedback influenced memory consolidation processes, we next examined performance levels of the surprise memory recognition test 24 hours later. About half of the participants performed a free recall test, while the other half performed a forced-choice recognition test (with 25% chance level, see Methods & Materials). Since under free-recall, participants’ performance was very low (22.1% overall and 4.4% on M+) floor effects are likely. In contrast, performances were substantially higher on the recognition test and comparable in size with our previous work using the same task and memory test (M+: 42.8%; M-: 36.5%; Ripollés et al., 2016, 2018), thus only the data from the recognition group is analyzed below.

To characterize how the presence of feedback affects recognition memory, we focused our analysis on the subset of new-words which were correctly answered on day 1, and could, thus, be either remembered or forgotten a day later in accord with our earlier approach (see Fig. 2b; Ripollés et al., 2016, 2018). Participants were, on average, more likely to remember the congruent new-words (congruent: 62.4% versus incongruent: 38.9%, main effect of congruence: χ^2^(1) = 78.32 *p* < .0001) after 24h. This pattern remained significant when comparing the raw recognition rates of congruent and incongruent trials (i.e. independent of whether they were correctly solved on day 1; χ^2^(1) = 9.18, p < .002) and is hence not due to the smaller number of to-be-remembered M+ items on day 1.

Importantly, the timing of presence of feedback (first or second half) significantly influenced the probability to remember, but differently so for DA-dependent and DA-independent learning: For congruent new-words only, withdrawing feedback lead to a significant drop in memory perfortmance (62.8% vs 50.4%, odds-ratio=1.66±0.4, *p* = .034) while the decrease in performance did not significantly differ for the group who started off without feedback (odds-ratio=0.75±0.18, *p* = .24; for congruent only, interaction of feedback by order: odds-ratio=2.21±0.75, *p* = .021). Post-hoc comparisons revealed that the probability to remember in the absence of feedback, significantly differed depending on whether the new-words were learned during the first or second half on the previous day (i.e. before or after participants started to receive feedback during learning; odds-ratio=0.42, *p =* .013), while there were no order differences for new-words which were learned while feedback was provided (*p* = .79; i.e., memory recognition was not affected if feedback was provided first or after the no-feedback condition).

In contrast, this pattern was absent for the new-words which had been associated with two incongruent sentences (3-way interaction of feedback by order by congruence: χ ^2^(1) = 4.74, *p* = .03). We observed a trend for feedback improving the memory for the incongruent new-words (for incongruent only, main effect of feedback: odds-ratio=1.62±0.45, *p* = .08), independently of when it was received (for incongruent only, interaction effect of feedback by order: odd-ratio=0.85±0.23, *p* = .55). The probability to correctly remember the incongruent new-words was also significantly above chance under all four conditions (all *p’s* < .044 bonferroni-corrected).

In sum, this pattern of results suggests that, during our task, intrinsic- or extrinsic-generated DA-dependent signals interact and affect recognition memory. Importantly, participants benefited less from external feedback after being exposed to a period of self-regulated learning (internal DP-dependent signals) without no externally regulated feedback. Overall pattern of responses on day 2 was markedly different from the pattern observed on day 1.

### Subjective evaluation of trial performance

Behavioral responses are often accompanied by particular reward-related emotional, motivational and metacognitive states. Previous studies on our new-word learning paradigm demonstrated that an increase in dopaminergic signaling was tightly linked to enhanced pleasure and confidence ratings, whereas arousal ratings were not enhanced (Ripolles, 2016, 2018). To evaluate the influence of dopaminergic signaling in the current study more directly, we also measured trial-based arousal and pleasantness ratings on day 1 in addition to performance. Replicating previous findings, for pleasure ratings we again found a larger difference for correct relative to incorrect responses in the no-feedback M+ conditions than the no-feedback M-conditions (see Supplemental Material 1,3). No such differences between no-feedback M+ and M-conditions were observed for the arousal ratings, again replicating previous observations (see Supplemental Material 2,4). Moreover, we also observed an effect of feedback on pleasantness ratings following correct vs. incorrect responses, with larger differences for feedback trials. Please note that ratings were obtained after feedback, hence this rating likely reflects dopaminergic signaling due to feedback in addition to intrinsic signaling. Significantly extending previous observations, we also observed a drop in pleasantness ratings for the feedback-first group after feedback was withdrawn (see Supplemental Material 1,3). Strikingly, this decrease in pleasantness ratings was not mirrored by online task performance on day 1, but by recognition performance on day 2.

In previous studies we had never assessed subjective states during the recognition period. Here, adding to previous observations, we also assessed subjective states on day 2; in particular arousal, pleasantness plus confidence (note that confidence was not included on day 1 since ratings were acquired after feedback; no feedback was provided on day 2 rendering the confidence judgements more meaningful). For pleasantness, we observed higher ratings for no-feedback M+ trials than feedback M+ trials and opposite effects for M-trials suggesting that external feedback on day 1 had interfered with intrinsic dopaminergic signaling (Supplemental Fig 5). Moreover, we observed higher pleasantness ratings for remembered M+ trials relative to incorrect and forgotten M+ trials; and opposite effects for M-trials suggesting enhanced dopaminergic signaling during correct remembering which may indicate additional dopaminergic reconsolidation processes (Redondo et al., 2011). Arousal ratings did not show any differences on day 2 (Supplemental Fig 6).

Finally, confidence ratings, which may reflect complementary metacognitive processing, were most enhanced for remembered M+ words relative to forgotten and incorrect M+ words in the no feedback first group. Additionally, we also observed robust effects for confidence ratings of incorrect M-words in this group (i.e. participants incorrectly assigned a meaning to M-words on day 2, although they had correctly indicated the absence of congruent meaning on day 1). These response characteristics (incorrect responses with high confidence) may indicate that participants erroneously recalled false memories. Together this pattern of results suggests that starting without feedback led to a self-centred strategy on day 1 which resulted in enhanced recognition on day 2 together with metacognitive awareness but at the cost of higher confidence in false memories (i.e. enhanced confidence ratings in the M-incorrect category, see suppl. Material p.14, SFig. 7 left bottom).

## Discussion

In this study we probed the interplay of external (extrinsic) and internal (intrinsic) motivational signals on recognition memory by manipulating the presence of informative feedback in the context of DA-dependent and DA-independent learning. We found that these signals had distinct effects on immediate task performance during learning (day 1) and on more durable consolidation processes as evidenced by memory performance on day 2. Importantly, we observed that the impact of an extrinsic reward (i.e. external feedback) on a DA-dependent memory depended on whether an intrinsically generated signal had been already established (i.e., on whether self-regulated learning, without feedback, was driven by an intrinsic reward signal tied to successful meaning extraction). The influence of DA on congruent trials was signified by elevated pleasure ratings for correct relative to incorrect M+ trials, while the difference in pleasure was smaller for M-trials. Moreover, withdrawal of feedback led to reduced pleasure ratings and a significant decrease in recognition on day 2 for DA-dependent M+ but not DA-independent M-learning. Also on day 2, pleasure ratings were enhanced for correctly remembered M+ words, while no effects of arousal were evident on day 2. Finally, confidence ratings, which may reflect additional metacognitive strategies, were enhanced for M+ words in the group which started without feedback as well as for false memories. Remarkably, the enhancement was independent of whether or not feedback was provided, suggesting more robust memory representations potentially caused by initially relying on self-regulated learning which may have lasted even after feedback was introduced but at the cost of higher confidence in wrongly remembered words with no congruent meaning.

During day 1, adding external feedback improved performance, especially if it was provided in the second half, that is, after an initial, self-guided learning period without any feedback. Remarkably, this effect did not depend on what was to be learned – a specific meaning extracted from two congruent sentences (DA-dependent) or simply the fact that the sentences were incongruent (DA-independent). In stark contrast, on the surprise memory test performed by the participants a day later we observed markedly different effects between the congruent and incongruent conditions. On day 1, participants more often correctly indicated that two sentences had been incongruent; on day 2, however, they were less likely to remember this fact about the new-words as compared to the likelihood of remembering a specific meaning of a new-word. This large difference between congruent and incongruent new-words on memory recognition could be attributed to the well-known memory congruency effect (Tse et al., 2007; van Kesteren, Ruiter, Fernández, & Henson, 2012; Greve et al., 2019). Incorporating new information into long-term memory that is congruent with previous knowledge or “scheme” is easier than assimilating incongruent information. In this sense, when successfully discovering the meaning of a new-word, the semantic congruency of the previous two sentences might help to integrate this new trace into a coherent knowledge prior or scheme outside the SN/VTA-HP loop that might in turn reinforce encoding and later retrieval. A knowledge prior might be represented in prefrontal structures most likely inferior frontal gyrus (see Ripolles, 2014, Fig. 4) or superior frontal regions.

The online performance on day 1 improved substantially more when feedback was introduced in the second half of the task (for both congruent and incongruent trials) revealing a time-dependent effect. Note that feedback itself could not directly enhance performance, because it was given after each judgement was made and the information that one had just made a mistake could not help to perform better on the subsequent trials. One possibility is that adding performance feedback boosts motivation, leading to more attentional and cognitive resources being deployed. However, then performance should be lower on trials without feedback in general, and not in this time-dependent manner. Alternatively, performance might be boosted in tasks whose structure (also referred to as “learning set”, Harlow, 1949) had been already learned in a self-guided manner before the introduction of the extrinsic incentives. Then, the additional informative feedback about successful task performance could further strengthen these already established processing routines, leading to an increased performance on day 1.

On day 2, participants remembered slightly less that a new-word lacked a congruent meaning for those words which had been presented in trials with feedback, in comparison with those presented without feedback). Thus, when learning occurred in the absence of an intrinsically triggered DA-signal (incongruent trials), external rewards in form of informative feedback improved performance on the same day, as well as moderately improved memory recognition a day later, potentially by adding the DA-signal needed for LTP. Similar results have been obtained, for example, for visual working memory (Adam and Vogel, 2016) and learning of a list of words (Nielson, Bryant, 2005). This can be seen in light of competence (Deci et al. 1999), as feedback helps learners master the task at hand by facilitating evaluation and monitoring of learning performance (Clark 2014; Rohlfing et al. 2016; Kuhl 2010), thereby strengthening self-regulation (Boekaerts 1999; Donker et al. 2014). Indeed, we observed at least a small enhancement for both memory and online performance in the presence of feedback. Concordantly, pleasantness ratings were also enhanced for correct trials if presented with feedback relative to incorrect trials and this effect was enhanced for both M+ and M-trials. In sum, our results are in line with the highly consistent main effect of extrinsic rewards (Cerasoli et al., 2014) on online performance but extends earlier observations by showing that the impact of feedback is dependent on the trial-specific DA-level.

In contrast to the incongruent trials, the results for DA-dependent learning (congruent trials) showed a more complex pattern. Specifically, the pattern of online performance (day 1) mimicked the one of the incongruent trials, such that receiving feedback in the second half improved task success. While withdrawing feedback in the second half did not lead to a change in the online performance of day 1, it profoundly affected recognition memory on day 2. Specifically, there was a drop in recognition rates of new-words which had to be learned without feedback after a preceding phase with feedback. This is akin to the well-known *undermining effect* (for a review, see Deci, Koestner, Ryan, 1999). Importantly, a drop in performance tied to the undermining effect has previously been related to a drop in activity in the VS and SN/VTA (Murayama et al., 2010). Since we observed this pattern only in DA-dependent (congruent) and not on DA-independent (incongruent) trials, our results strongly suggest that a potential factor determining whether an undermining effect will be observed is the involvement of the SN/VTA-HP loop and the associated dopamine release (Ripollés et al., 2016, 2018).

A possible alternative explanation for this feedback-order dependence is the increasing levels of fatigue over the course of the experiment. Hence, with more trials more fatigue is present and adding feedback in the second half could counteract this fatigue-based effect. However, it is unclear why fatigue would affect DA-dependent learning more than DA-independent learning. Instead, learning the task structure (Harlow, 1949) in a self-regulated manner might be accompanied by a more robust, intrinsically generated DA release than learning the task with external feedback and the associated extrinsic DA release. From this perspective, adding external feedback in the second half leads to a slight decrease in memory, potentially due to the interaction of external and internal DA-signaling, while removal of the feedback leads to the removal of the main source of the DA signal (the feedback) while intrinsically guided DA-release is not established and, thus, prevents DA-dependent LTP from taking place. Accordingly, overall pleasure dropped when feedback was withdrawn. Arousal ratings dropped as well. However, there was a difference between correct and incorrect M+ and M-trials for pleasure ratings but not for arousal ratings, again suggesting an amelioration of DA-release. This would explain why the undermining effect is only observed in the DA-dependent (congruent) trials. Future neuroimaging work should be able to address this question.

Our results for the congruent trials are also in line with previous research proposing that insight (i.e., a sudden realisation of something, in our paradigm, the meaning of the word to be learned) improves long-term memory (Auble et al., 1979, Domonowski and Buyer, 2000; Ash et al., 2012; Danek et al., 2013; Kizilirmak et al., 2016a, 2016b). Moreover, Murayama and Kuhbander (2011) showed that subsequent memory traces for answers to trivia questions were improved by providing monetary reward for correctly answered trivia questions and that this effect was driven by the uninteresting questions. This suggests that the more interesting trivia questions trigger the SN/VTA-HP loop on their own (Gruber et al., 2014), similar to our congruent sentences and that the less interesting ones are DA-independent, akin to our incongruent trial, leading to the observed pattern. In line with our results, Murayama and Kuhbander (2011) also observed a stark difference when comparing the effects of monetary reward between immediate and delayed memory tests (with the latter showing significant improvement). Moreover, in their quantitative meta-analysis, Cerasoli and colleagues (2014) found that different measures of performance (quantitative versus qualitative) show a differential link to intrinsic motivation and extrinsic rewards. They suggested that this might be due to those metrics being used for different types of tasks – quantitative ones for more boring and repetitive tasks, and qualitative metrics for tasks involving creativity and learning. Overall, this suggests that the effects of reward on immediate performance and memory consolidation processes can differ systematically. We propose that these distinct patterns might be caused by the involvement (or absence) of dopaminergic long-term potentiation processes, in line with the hippocampus-dependent memory consolidation model (Adcock et al., 2006; Lisman and Grace, 2005).

In addition, confidence in correctly remembered words and false memories were also elevated in participants based on the timing of feedback. Likely, starting without feedback led to a longer-lasting strategy to rely on self-monitoring. Confidence in memory and its related metacognitive processes have previously been linked to activation in the cingulate cortex (Moritz et al., 2006), but also to activation in striatal reward areas (Molenberghs et al., 2016). Striatal reward regions are also active during our task especially during the successful meaning extraction of congruent trials (Ripolles, 2014, 2016). In addition to enhanced confidence for correctly remembered items these ratings were also enhanced for false memories. The formation of false memories, however, was not reflected in our arousal or pleasantness ratings on day 1 or day 2 suggesting an alternative origin of this effect after initial encoding.

These results have important implications for language learning in educational contexts as it provides a mechanistic explanatation for the benefits of self-regulated learning and intrinsic motivation; concepts which are implemented in Montessori schools among others (c.f. Paris, Paris, 2001; Clark, 2012). They also have implications regarding clinical populations, as earlier studies with aphasic patients (Peñaloza et al. 2016, 2017) suggest that informative feedback enhances performance. In this vein, our results suggest that clinical patients (given that brain areas processing feedback are not too impaired; Stone et al. 2002; Haber and Knutson 2009; Murayama et al. 2010; Vrtička et al. 2014) engaging in DA-independent learning should profit more from paradigms in which feedback is provided. In contrast, our data do not support the notion that immediate task performance (unlike longer-term memory) will always deteriorate when feedback starts to be withheld (Carr and Walton, 2014).

In sum, our work shows that intrinsic and extrinsic reward signals interact in a time-dependent manner and the precise pattern of the ensuing effects depends on whether they involve dopaminergic memory consolidation processes. Our results highlight the distinction between online performance and consolidation-related memory retrieval processes and extend our understanding of how intrinsic motivation and extrinsic incentives interact by taking the DA-dependent and DA-independent context into account. Moreover, memory encoding was also accompanied by enhanced pleasure ratings for later remembered words and starting without feedback led to enhanced confidence during retrieval for both right and wrong memories. Together, this work suggests that future educational programs should incorporate self-regulated learning without feedback when intrinsic DA-dependent learning is bound to occur, but also highlights the risks of having higher confidence in false memories during self-regulated learning.

## Methods & Materials

### Participants

A group of sixty healthy, right-handed native German speakers took part in the experiment (*M*_age_ = 24.27 years, *SD*_age_ = 3.8, 40 females). Participants were recruited on campus of the Otto-von-Guericke University using emailing lists and poster announcements. Inclusion criteria were normal or corrected to normal vision, German as first language, and no self-reported history of mental illness. The study was carried out in accordance with local ethics and all volunteers gave their written informed consent to participation at the beginning of the experiment.

The original sample size was chosen to be approx. 30 participants per group (recall / recognition memory test, see below). This sample size was selected based on several criteria, including the recommendation that, in order to achieve 80% of power, at least 30 participants should be included in an experiment in which the expected effect size is medium to large (Cohen, 1988). In addition, we took into account the sample sizes of previous studies using this paradigm which produced significant effects (range: between 24 and 40 participants; Ripollés et al., 2014, 2016, 2018; Angwin et al., 2019). To account for potential drop-out due to the experiment spanning two consecutive days, we aimed to recruit 33 participants per group (based on previous experience in our lab of a 10% attrition rate). The final sample size was 31 participants in the free-recall group and 29 in the recognition test group (see below). Further mixed models were used for statistical analyses to maximise sensitivity (Gelman, Hill, 2006).

### Materials

This study uses the same paradigm as our previous work (Ripollés et al., 2014; 2016; 2018), except that we added performance-contingent feedback (Deci et al. 1999; Ryan et al. 1983) on half the trials. Stimuli were presented using Psychtoolbox (http://www.psychtoolbox.org) in MATLAB R2012b (The MathWorks, Inc., Natick, MA, USA).

Feedback consisted of either happy or sad smileys, indicating correct or incorrect responses respectively (see Fig. 1). To control for the perception of faces (Haxby et al., 1999; Puce et al., 1999) and to match the trial duration, phase transformed images of smileys were used (Allison et al., 1999; Liu et al., 2002) in the no-feedback condition. Further, to prevent attributing a positive or negative meaning to any single phase-transformed stimulus, each scrambled feedback stimulus was displayed only once (Fig. 1; for more details on the materials & procedure see Supplementary Material).

### Procedure

As in our previous work, the experiment consisted of a learning phase (approx. 2h) and a memory-test phase (approx. 30min) a day later (M=23h, SD = 50min).

### Learning session

On the first day, participants read 80 pairs of sentences which ended in a new-word. Importantly, these pseudo-words were presented as part of either two congruent (40 pairs) or two incongruent sentences (40 pairs). In the congruent condition, a single meaning of a pseudo-word could be inferred from the two sentences (e.g., 1 “Every Sunday the grandmother went to the *jedin*” and 2. “The man was buried in the *jedin*”; here, “jedin” means graveyard and is congruent with both the first and second sentences). For incongruent sentence pairs no single coherent meaning could be inferred (e.g. 1. “Every night the astronomer watched the *heutil*” here moon is a possible meaning of “heutil; but 2. “In the morning break co-workers drink *heutil*”; here, coffee is now a possible meaning which is inconsistent with the first sentence).

In order to mimic real-life contextual learning where novel words can be encountered at different times and in different contexts, first and second sentences ending in the same new-word, were presented separated in time and in sets of eight pairs (see Fig. 1a). In ten different blocks, participants were presented with four congruent and four incongruent pairs of sentences. In particular, participants first read all eight first sentences, each paired with a novel new-word; then, they read the eight second sentences paired with the same new-words which were presented in randomized order relative to the first presentation. Immediately after finishing reading each second sentence and the associated new-word, participants were asked to verbally indicate the meaning of this new-word. This response was coded manually as correct/incorrect by the experimenter. Crucially, in half of the blocks, participants received infomative feedback, and in the other half no-informative feedback (Fig.1b). The order was counter-balanced across participants (feedback in 1st half / feedback in 2nd half). After the feedback screen, participants further rated their subjective state, along pleasantness and arousal using a 9-point Likert scale semantic differential. For pleasantness, participants rated how they pleasant they were feeling at this momemt, and for arousal how calm they were. On day 1 scales were given after the feedback to ensure that participants linked feedback to the meaning task and not to their subjevctive ratings which could have happened if we had asked for self-evalutaions prior to feedback.

### Surprise Memory test

Participants were told that the session on the second day would involve a personality evaluation, which would enable the investigation of the role of individual differences in personality on mood changes during reading. Instead, participants had to take a surprise memory test. 31 participants underwent a cued free-recall memory test, while the remaining 29 participants took a recognition test.

In the free-recall task, participants were presented with all 80 new-words from the learning session and had to verbally indicate what its meaning was (for M+ new-words), that it had been paired with incongruent sentences (for M-new-words), or that they were unsure (“I don’t know”). Their responses were manually coded by the experimenter. Participants were uncertain for the majority of the new-words (53.8% of trials), and answered correctly only 18.2% of the trials which were mostly the incongruent trials – only 4.4% of the congruent new-words were correctly recalled on the seconds day (i.e. between 1 and 2 new-words on average with only 12 participants showing any memory trace for any of the 40 M+ words at all). Due to this low performance, floor effects are extremely likely and we did not analyze the free-recall data of day 2 any further.

In the recognition test, participants were presented with all 80 new-words from the learning session in random order and without their respective sentences or feedback, while given four response options: 1) the meaning evoked by the second sentence in which the new-word had been presented (i.e. the correct option for congruent trials, but a lure for incongruent trials), 2) with a lure (a randomly selected meaning from the remaining new-words presented during the learning phase), 3) an “incongruent” option (i.e., to signal that the new-word had no congruent meaning), and 4) an “I don’t know” response option to indicate that they were uncertain.

Additionally, participants were asked to provide pleasantness and arousal ratings after each word (using the same scales as on day 1). Further, participants rated their confidence that their answer had been correct (I am uncertain vs. certain) on a 9-Likert scale. All scales were poled in the same direction, with low ratings reflecting lower pleasantness, confidence and higher arousal

### Statistical analysis

All statistical analyses were conducted in R (version 3.6.1). To assess the performance during the learning session, we estimated a generalized linear mixed model with a logistic link function, for either correct or incorrect responses. This model is conceptually comparable to a logistic regression, but explicitly models the repeated measurement of responses from the same participant in multiple conditions (e.g. Gelman, Hill, 2007). Specifically, to account for the repeated-measures nature of the dataset we specified a random intercept per subject as a random factor. In addition, we included as fixed effects the main factors of interest, namely congruence (congruent/incongruent), feedback (feedback/no-feedback), and order (feedback first/no-feedback first), as well as all their interactions.

Analogously, for the recognition test we estimated a logistic mixed model for either remembering or forgetting the meaning of a new-word (i.e. correctly indicating a specific meaning for congruent trials and correctly answering “incongruent” for the incongruent trials), with the same fixed and random effects structure.

For model estimation, we used the *mixed* function in the *afex* package (version 0.18; Singmann, et al., 2016) which in turn uses the *lme4* package (version 1.1; Bates, et al., 2015) for the estimation of the models; *p*-values were computed using the Satterthwaite approximation of the degrees of freedom when assessing the significance of the fixed effects, as implemented by the *mixed* function. Importantly, mixed models account for different numbers of participants per factor level and still provide an unbiased estimate of effects (Gelman, Hill, 2007). Post-hoc comparisons were calculated using the *emmeans* package (Lenth, 2019).

See supplementary materials for the statistical analysis of the subjective ratings.

## Data availability statement

The behavioral data that support the findings of this study, as well as the analysis code are available via OSF with the identifier doi: *[final DOI will be generated after review to include also any potential reviewer-requested analyses]*

## Notes on supplementary analyses

Incorrect responses including erroneous and “I don’t know” responses. On day 1, participants answered “I don’t know” on 124 trials out of 4800 (2.58%). On day 2, participants answered 1884 out of 4800 (39.25%), when considering both free recall and recognition together; for the recognition test alone, they answered “I don’t know” 549 out of 2320 trials (23.66%). Even though potentially being driven by slightly different processes, we pool the incorrectly answered trials with those where participants indicated “I don’t know” for these subsequent analyses. Note that we did not assess trial-based confidence on day 1; thus, a further grouping of “I don’t know” responses was not possible.

To consider the ratings on day 1, we report two sets of analyses:

1. All 60 participants while considering their learning performance, but independent of their subsequent memory performance on the next day.
2. Only the participants who did the recognition test on day 2, while splitting the ratings according to the combination of day 1 and day2 (i.e. remembered vs forgotten vs incorrect).

## Pleasantness on day 1 ratings based on learning performance

**Supplemental Fig. 1:**
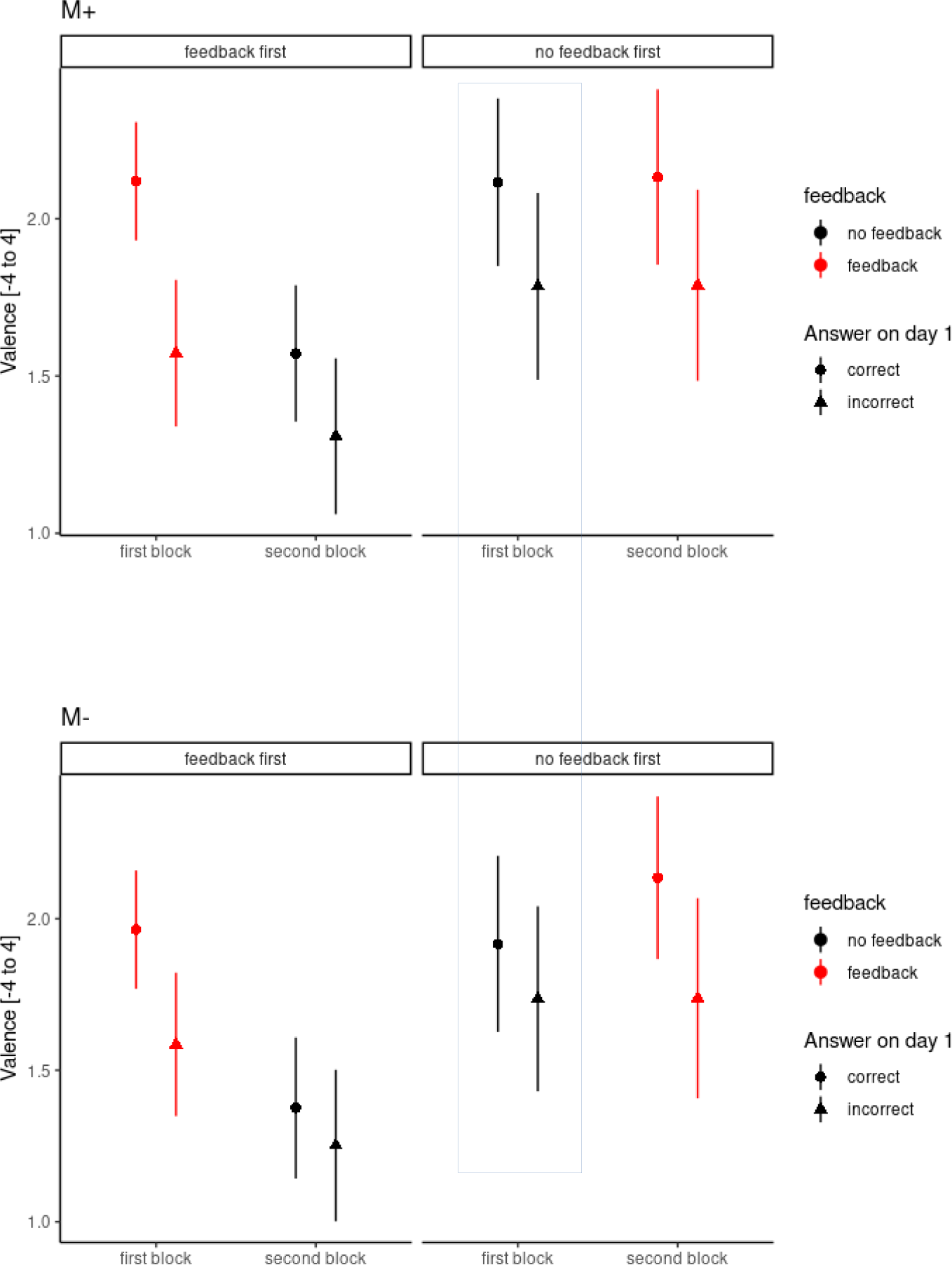
Condition-specific group means and between-subject SEM for pleasantness ratings on day 1 sorted by performance on day 1 (correct/incorrect). Upper panel: M+ (dopamine-dependent trials). Lower panel: M-(dopamine independent trials). Left part of graphs: feedback first group; Right Part of graphs: no feedback first group. Framed area highlights the conditions similar to Ripolles et al., 2016.

**Supplemental Table 1:**
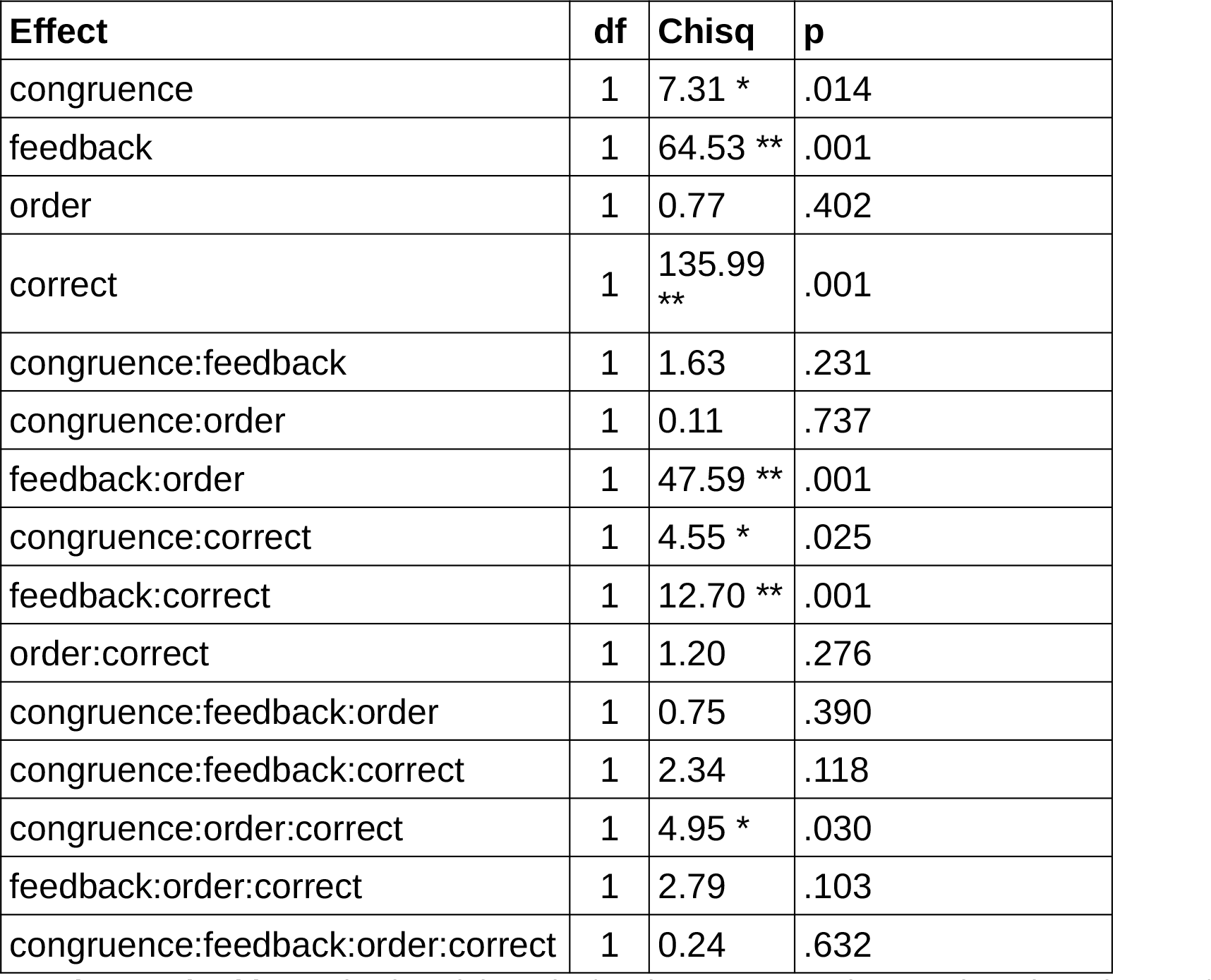
Mixed model results for pleasantness ratings on day 1 based on performance on the same day. The mixed model was estimated with *congruence*, *feedback*, *feedback-order* and *correct/incorrect* as independent variables. A random intercept per subject was included. P-values were calculated with parametric bootstrapping. df = degrees of freedom; Chisq = chi square value; p = probability. “X:Y” denotes the interaction of X with Y.

| Effect | df | Chisq | p |
| --- | --- | --- | --- |
| congruence | 1 | 7.31 * | .014 |
| feedback | 1 | 64.53 ** | .001 |
| order | 1 | 0.77 | .402 |
| correct | 1 | 135.99 ** | .001 |
| congruence:feedback | 1 | 1.63 | .231 |
| congruence:order | 1 | 0.11 | .737 |
| feedback:order | 1 | 47.59 ** | .001 |
| congruence:correct | 1 | 4.55 * | .025 |
| feedback:correct | 1 | 12.70 ** | .001 |
| order:correct | 1 | 1.20 | .276 |
| congruence:feedback:order | 1 | 0.75 | .390 |
| congruence:feedback:correct | 1 | 2.34 | .118 |
| congruence:order:correct | 1 | 4.95 * | .030 |
| feedback:order:correct | 1 | 2.79 | .103 |
| congruence:feedback:order:correct | 1 | 0.24 | .632 |

Pleasantness ratings on day 1 revealed interactions of *Feedback* with *Order* and with *Correctness* in addition to the main effects of *Congruence*, *Feedback* and *Correctness*. Moreover, trial *Congruence* interacted with *Order* and *Correctness* (triple interaction). The interaction of *Feedback* with *Order* indicated that a drop in pleasantness ratings occurred if feedback was withdrawn whereas it slighthly increased ratings if feedback was added later. These effects were independent of *Correctness* or *Order*. Additionally, differences in pleasantness ratings between correct and incorrect trials where larger for the congruent than incongruent trials. While this effect might be significant, it should be interpreted with caution since pleasantness ratings were obtained after performance feedback. Thus, the ratings after feedback may be more influenced by the feedback than by the subjective evaluation of task performance. Likewise, the difference in pleasantness rating for correct and incorrect trials were slightly higher for congruent trials in the group starting with feedback, whereas the difference was slightly higher for M-trials in the group starting without feedback. Note again, that this comparison collapses across no-feedback and feedback trials, thus might be difficult to interpret as it may contain influences from processes related to trial performance and feedback in some trials but not in others. Finally, we directly tested for a replication of the effect of pleasantness for the no-feedback trials (larger difference for correct-incorrect trials in the M+ but not the M-condition; Ripolles et al., 2014,2016,2018). The descriptive statistics indicate that pleasantness ratings were highest for the M+ correct trials (see SFig.1; framed region) despite the longer delay between response and pleasantness ratings than in our previous studies (in the framed region, all pairwise comparisons with M+ correct trials: *p*s < .014). This effect was also observed for the no-feedback trials of the feedback first group

## Arousal on day 1 ratings based on day 1 performance

Errorbars: between-subject SEM

**Supplemental Fig. 2:**
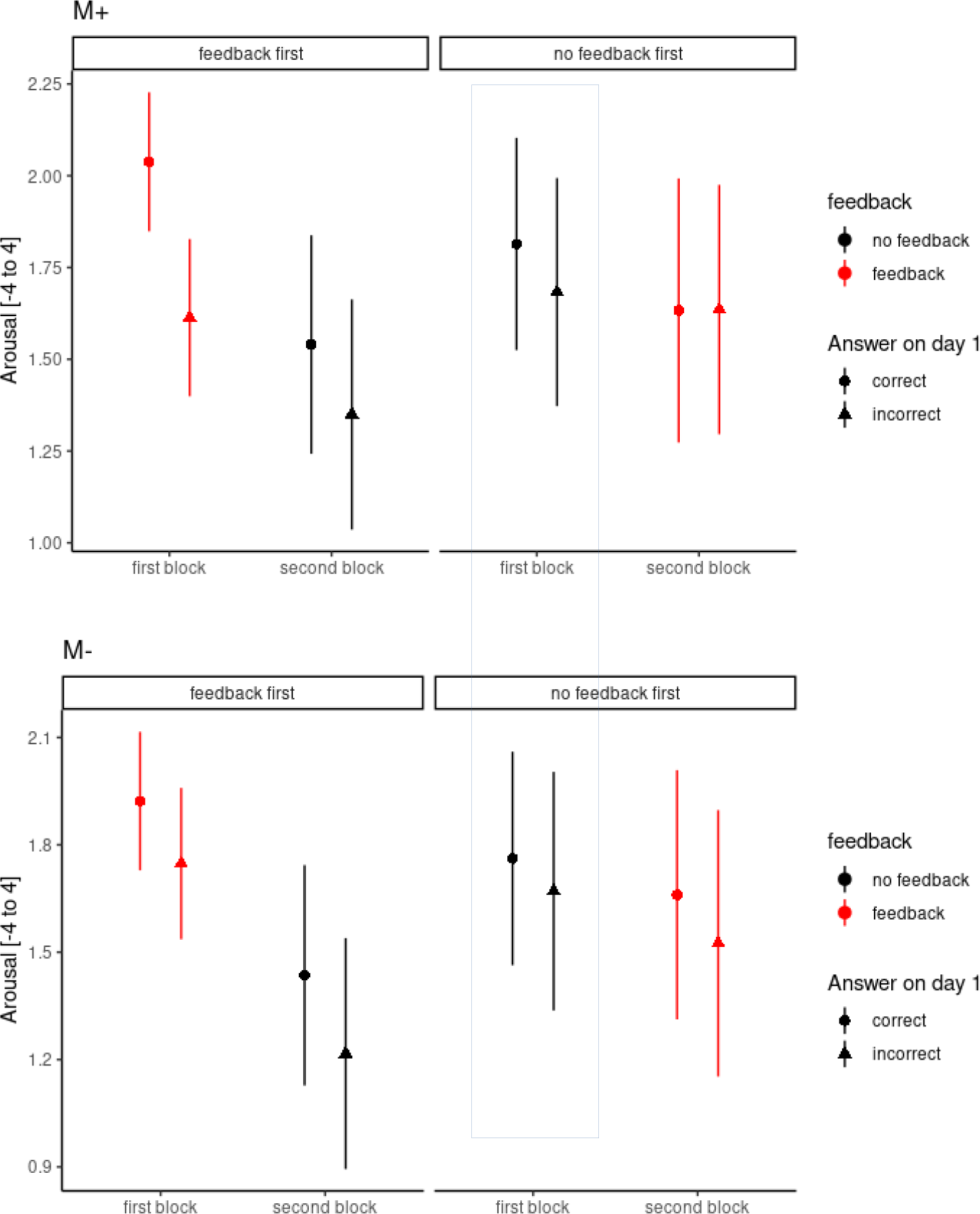
Condition-specific group means and between-subject SEM for arousal ratings on day 1 sorted by performance on day 1 (correct/incorrect). Upper panel: M+ (dopamine-dependent trials). Lower panel: M- (dopamine independent trials). Left part of graphs: feedback first group; Right Part of graphs : no feedback first group. Framed area highlights the conditions similar to Ripolles et al., 2016.

**Supplemental Table 2:** Mixed model statistical results for arousal ratings on day 1 based on performance on the same day. Mixed models were estimated with *congruence*, *feedback*, *feedback-orde*r and *correct/incorrect* as independent variables. A random intercept per subject was included. P-values were calculated with parametric bootstrapping. df = degrees of freedom; Chisq = chi square value; p = probability. “X:Y” denotes the interaction of X with Y.

| Effect | df | Chisq | p |
| --- | --- | --- | --- |
| congruence | 1 | 1.54 | .221 |
| feedback | 1 | 20.83 ** | .001 |
| order | 1 | 0.01 | .915 |
| correct | 1 | 38.63 ** | .001 |
| congruence:feedback | 1 | 1.27 | .268 |
| congruence:order | 1 | 0.34 | .554 |
| feedback:order | 1 | 67.57 ** | .001 |
| congruence:correct | 1 | 1.58 | .215 |
| feedback:correct | 1 | 0.03 | .851 |
| order:correct | 1 | 1.45 | .236 |
| congruence:feedback:order | 1 | 0.35 | .561 |
| congruence:feedback:correct | 1 | 0.00 | .962 |
| congruence:order:correct | 1 | 4.17 * | .042 |
| feedback:order:correct | 1 | 4.99 * | .022 |
| congruence:feedback:order:correct | 1 | 0.08 | .779 |

In addition to two main effects of Feedback and Correctness we observed an interaction of Feedback with order, and a triple interaction of Feedback with Order and Correctness plus again a triple interaction of Congruence with Order and correctness (see STab 2 for details). The interaction of Feedback with Order was again caused by a drop in arousal ratings if feedback was withdrawn; further, arousal ratings to correct relative to incorrect trials were most pronounced in the group starting with feedback but only if feedback was provided yielding a significant triple interaction of Feedback with Order and Correctness. Moreover, the interaction of Congruence with Order and Correctness was driven by the difference in arousal ratings for correct vs. incorrect trials in M+ condition for the group starting with feedback. Finally, we directly tested for a replication of the null effect of arousal for the no-feedback trials (Ripolles et al., 2016,2018). The descriptive statistics indicate that arousal ratings were similar for the M+/M-correct trials (see SFig.2; framed region) resulting in no significant differences in arousal (main effect of congruence: p > .22, interaction of congruence and correctness: p > .78).

## Pleasantness on day 1 based on remembered/forgotten

**Supplemental Fig. 3:**
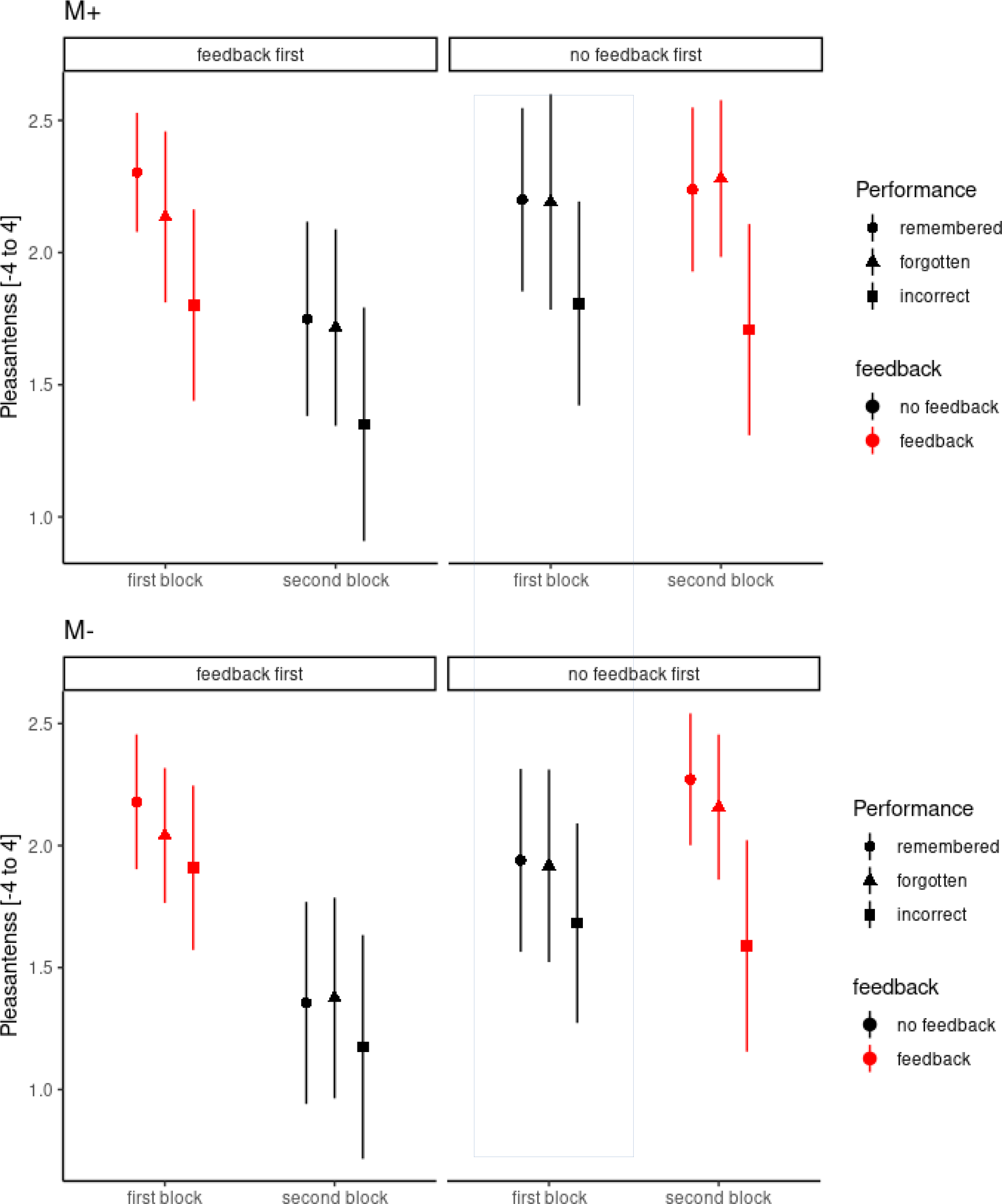
Condition-specific group means and between-subject SEM for pleasantness ratings on day 1 sorted by performance on day 2 (remembered/forgotten/incorrect). Upper panel: M+ (dopamine-dependent trials). Lower panel: M- (dopamine independent trials). Left part of graphs: feedback first group; Right Part of graphs : no feedback first group. Framed area highlights the conditions similar to Ripolles et al., 2016.

**Supplemental Table 3:** Mixed ANOVA statistical results for pleasantness ratings on day 1 based on performance on day 2 (remembered/forgotten). Mixed models were estimated with *congruence*, *feedback*, *feedback-order* and *rememForgot* as independent variables. Note that Factor “*rememForgot*” has 3 levels: Remembered (correct on both days), forgotten (correct only on day 1) or incorrect (already on day1 incorrect). A random intercept per subject was included. P-values were calculated with parametric bootstrapping. df = degrees of freedom; Chisq = chi square value; p = probability. “X:Y” denotes the interaction of X with Y.

| Effect | df | Chisq | p |
| --- | --- | --- | --- |
| congruence | 1 | 5.58 * | .018 |
| feedback | 1 | 62.16 ** | .001 |
| order | 1 | 0.18 | .680 |
| rememForgot | 2 | 56.18 ** | .001 |
| congruence:feedback | 1 | 5.81 * | .013 |
| congruence:order | 1 | 0.10 | .751 |
| feedback:order | 1 | 48.38 ** | .001 |
| congruence:rememForgot | 2 | 3.48 | .180 |
| feedback:rememForgot | 2 | 0.70 | .736 |
| order:rememForgot | 2 | 5.31 + | .073 |
| congruence:feedback:order | 1 | 0.00 | .951 |
| congruence:feedback:rememForgot | 2 | 3.20 | .217 |
| congruence:order:rememForgot | 2 | 3.23 | .217 |
| feedback:order:rememForgot | 2 | 1.43 | .503 |
| congruence:feedback:order:rememForgot | 2 | 1.85 | .402 |

We further assessed the relationship of subjective ratings with memory performance by grouping the correct trials into those that were remembered on day 2 vs. those that were forgotten (note that this was only done for the group of subjects who performed the recognition task (n=29), as free recall performance was too low). For pleasantness, we observed that pleasantness ratings were in between remembered and incorrect trials in almost all conditions (significant main effect of *Remembered/Forgotten*; see STab.3 plus SFig.3). Significant interactions were found for the factors *Feedback* with *Congruence*, *Feedback* with *Order* plus a trend for *Order* with *Remembered/Forgotten* (see STab.3). The interaction of *Feedback* with *Order* was virtually identical to the one described above (drop in performance after feedback was withdrawn), whereas the interaction of *Feedback* with *Congruence* was due the ratings after feedback being equal for M+ trials and M-trials (both on average 2.05), while ratings in absence of feedback differed between M+ trials (on averge 1.82) and M-trials (1.62, difference: *p* = .004). In addition, the trend towards *Order* and *Remembered/Forgotten* was caused by higher ratings for forgotten and remembered in the no feedback first group, as compared to the feedback first group. Finally, the pre-planned comparison of remembered vs. incorrect pleasantness ratings again indicated a larger difference for M+ than M-trials (*p* = .034).

## Arousal on day 1 based on remembered/forgotten

**Supplemental Fig. 4:**
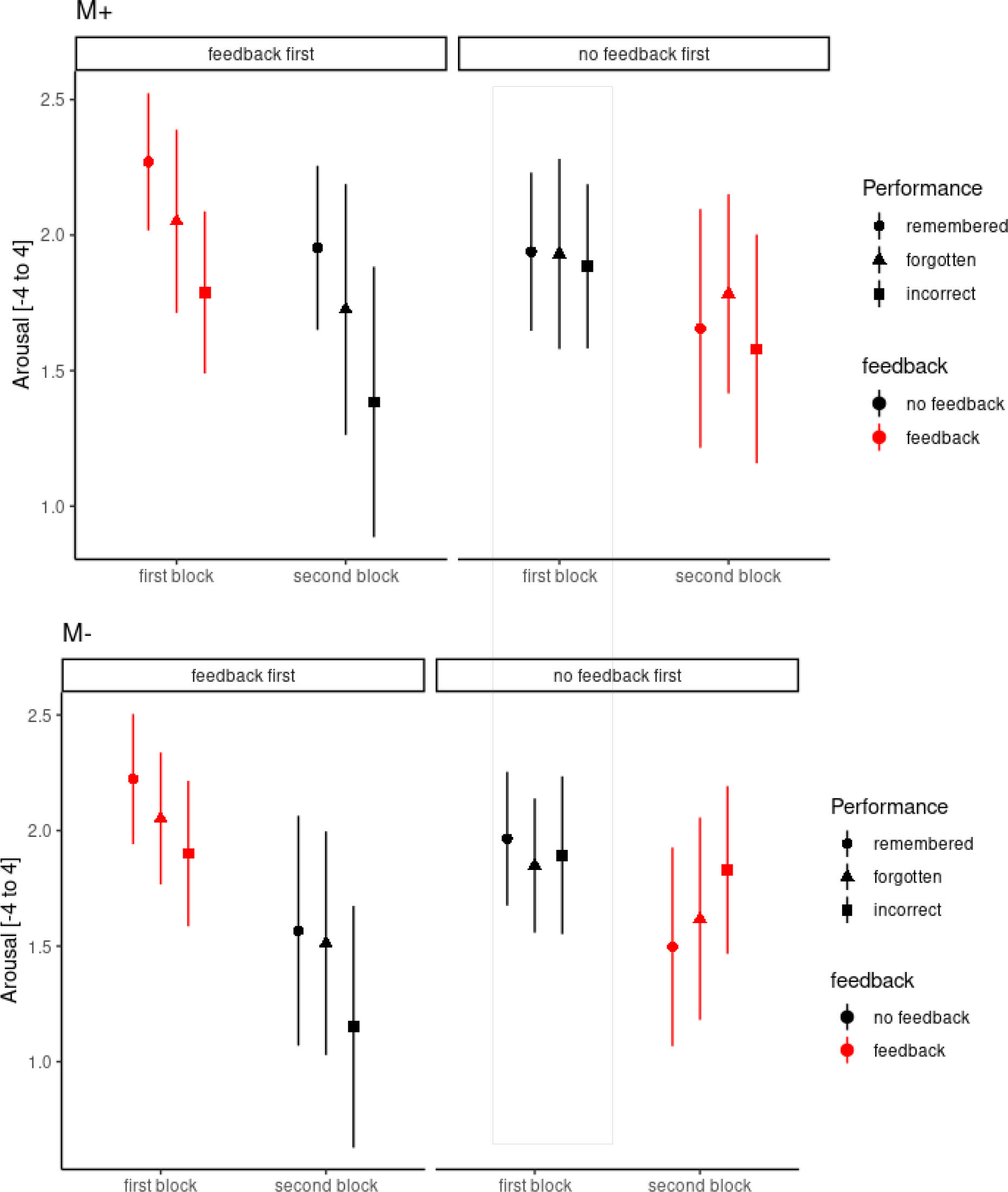
Condition-specific group means and between-subject SEM for arousal ratings on day 1 sorted by performance on day 2 (remembered/forgotten/incorrect). Upper panel: M+ (dopamine-dependent trials). Lower panel: M- (dopamine independent trials). Left part of graphs: feedback first group; Right Part of graphs : no feedback first group. Framed area highlights the conditions similar to Ripolles et al., 2016.

**Supplemental Table 4:** Mixed ANOVA statistical results for arousal ratings on day 1 based on performance on day 2 (remembered/forgotten). Mixed models were estimated with *congruence*, *feedback*, *feedback-order* and *rememForgot* as independent variables. Note that Factor “*rememForgot*” has 3 levels: Remembered (correct on both days), forgotten (correct only on day 1) or incorrect (already on day1 incorrect). A random intercept per subject was included. P-values were calculated with parametric bootstrapping. df = degrees of freedom; Chisq = chi square value; p = probability. “X:Y” denotes the interaction of X with Y.

| Effect | df | Chisq | p |
| --- | --- | --- | --- |
| congruence | 1 | 0.60 | .424 |
| feedback | 1 | 7.27 ** | .002 |
| order | 1 | 0.02 | .899 |
| rememForgot | 2 | 9.08 * | .013 |
| congruence:feedback | 1 | 1.52 | .212 |
| congruence:order | 1 | 0.18 | .663 |
| feedback:order | 1 | 69.87 ** | .001 |
| congruence:rememForgot | 2 | 2.74 | .267 |
| feedback:rememForgot | 2 | 15.27 ** | .001 |
| order:rememForgot | 2 | 5.04 + | .083 |
| congruence:feedback:order | 1 | 0.36 | .564 |
| congruence:feedback:rememForgot | 2 | 2.47 | .299 |
| congruence:order:rememForgot | 2 | 0.85 | .656 |
| feedback:order:rememForgot | 2 | 0.85 | .662 |
| congruence:feedback:order:rememForgot | 2 | 3.78 | .164 |

For arousal ratings on day 1 sorted by performance on day 2 we observed significant main effects of *Feedback* and *RememForgot* plus their interaction and the interactions of *Feedback* with *Order* and *rememForgot* with *Order*. The interaction of *Feedback* with *Order* was again due to the higher arousal ratings in the feedback first group when feedback was provided and lower arousal ratings without feedback relative to the no-feedback group. The interaction of *rememForgot with Order* was due to the monotonic increase in arousal ratings for incorrect to remembered trials in the feedback first group, which was not present in the no-feedback first group. Moreover, the interaction of *Feedback* with *RememForgot* was caused by the incorrect and forgotten trials having lower ratings in the absence of feedback (incorrect: difference 0.30, *p* < .001; forgotten: difference = 0.16, *p* = .045) as compared to those on trials with feedback, while the remembered trials did not differ significantly across feedback conditions ( *p = .14)..* Finally, the pre-planned comparison of remembered vs. incorrect arousal ratings again indicated no difference for M+ compared to M-trials (p>.63).

## Pleasantness on day 2 based on remembered/forgotten

**Supplemental Fig. 5:**
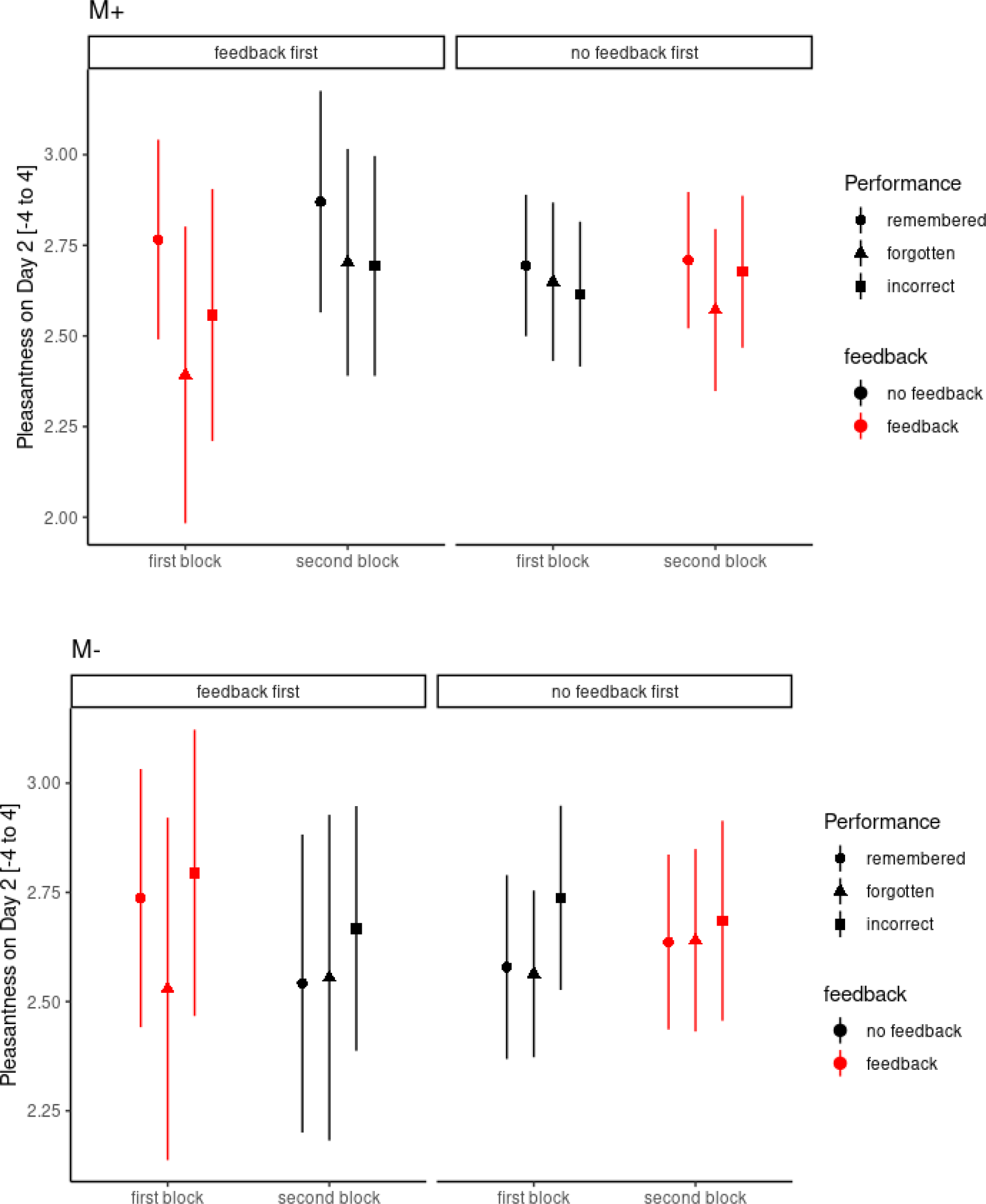
Condition-specific group means and between-subject SEM for arousal ratings on day 2 sorted by performance on day 2 (remembered/forgotten/incorrect). Upper panel: M+ (dopamine-dependent trials). Lower panel: M- (dopamine independent trials). Left part of graphs: feedback first group; Right Part of graphs : no feedback first group.

**Supplemental Table 5:** Mixed ANOVA statistical results for pleasantness ratings on day 2 based on performance on day 2 (remembered/forgotten). Mixed models were estimated with *congruence*, *feedback*, *feedback-order* and *rememForgot* as independent variables. Note that Factor “*rememForgot*” has 3 levels: Remembered (correct on both days), forgotten (correct only on day 1) or incorrect (already on day1 incorrect). A random intercept per subject was included. P-values were calculated with parametric bootstrapping. df = degrees of freedom; Chisq = chi square value; p = probability. “X:Y” denotes the interaction of X with Y.

| Effect | df | Chisq | p |
| --- | --- | --- | --- |
| congruence | 1 | 3.91 + | .053 |
| feedback | 1 | 0.00 | .953 |
| order | 1 | 0.00 | .978 |
| rememForgot | 2 | 5.70 * | .045 |
| congruence:feedback | 1 | 3.79 * | .046 |
| congruence:order | 1 | 1.76 | .199 |
| feedback:order | 1 | 0.69 | .405 |
| congruence:rememForgot | 2 | 16.09 ** | .001 |
| feedback:rememForgot | 2 | 0.93 | .638 |
| order:rememForgot | 2 | 1.08 | .549 |
| congruence:feedback:order | 1 | 2.58 | .113 |
| congruence:feedback:rememForgot | 2 | 0.44 | .834 |
| congruence:order:rememForgot | 2 | 3.82 | .158 |
| feedback:order:rememForgot | 2 | 1.51 | .479 |
| congruence:feedback:order:rememForgot | 2 | 0.48 | .788 |

For pleasantness ratings on day 2 we found a significant main effect of *rememForgot* plus a trend for *congruence,* their significant interaction (*congruence* x *rememForgot*), and the interaction of *congruence* with *feedback*. The interaction of congruence with feedback was due to higher pleasantness ratings for M+ trials without feedback but higher pleasantness ratings for M-trials with feedback on day 1 (Note that no further feedback was provided on day 2). The interaction of *congruence* with *rememForgot* was reflected in highest pleasantness ratings for remembered M+ trials relative to forgotten and incorrect trails and lower ratings for remembered and incorrect M-trials relative to incorrect trials. This pattern of results is similar to the pattern observed by Ripolles et al., 2016.

## Arousal on day 2 based on remembered/forgotten

**Supplemental Fig. 6:**
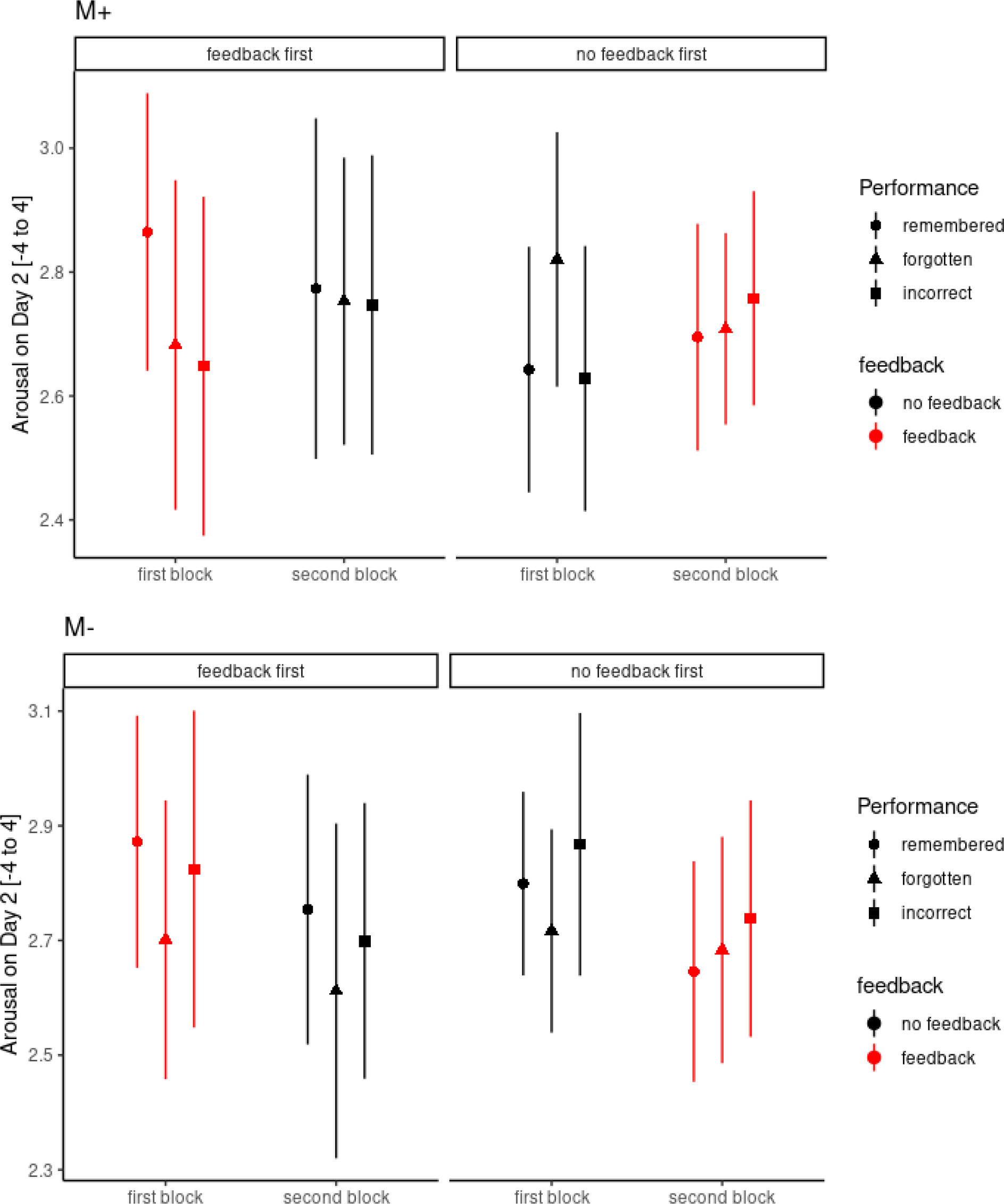
Condition-specific group means and between-subject SEM for arousal ratings on day 2 sorted by performance on day 2 (remembered/forgotten/incorrect). Upper panel: M+ (dopamine-dependent trials). Lower panel: M- (dopamine independent trials). Left part of graphs: feedback first group; Right Part of graphs : no feedback first group.

**Supplemental Table 6:**
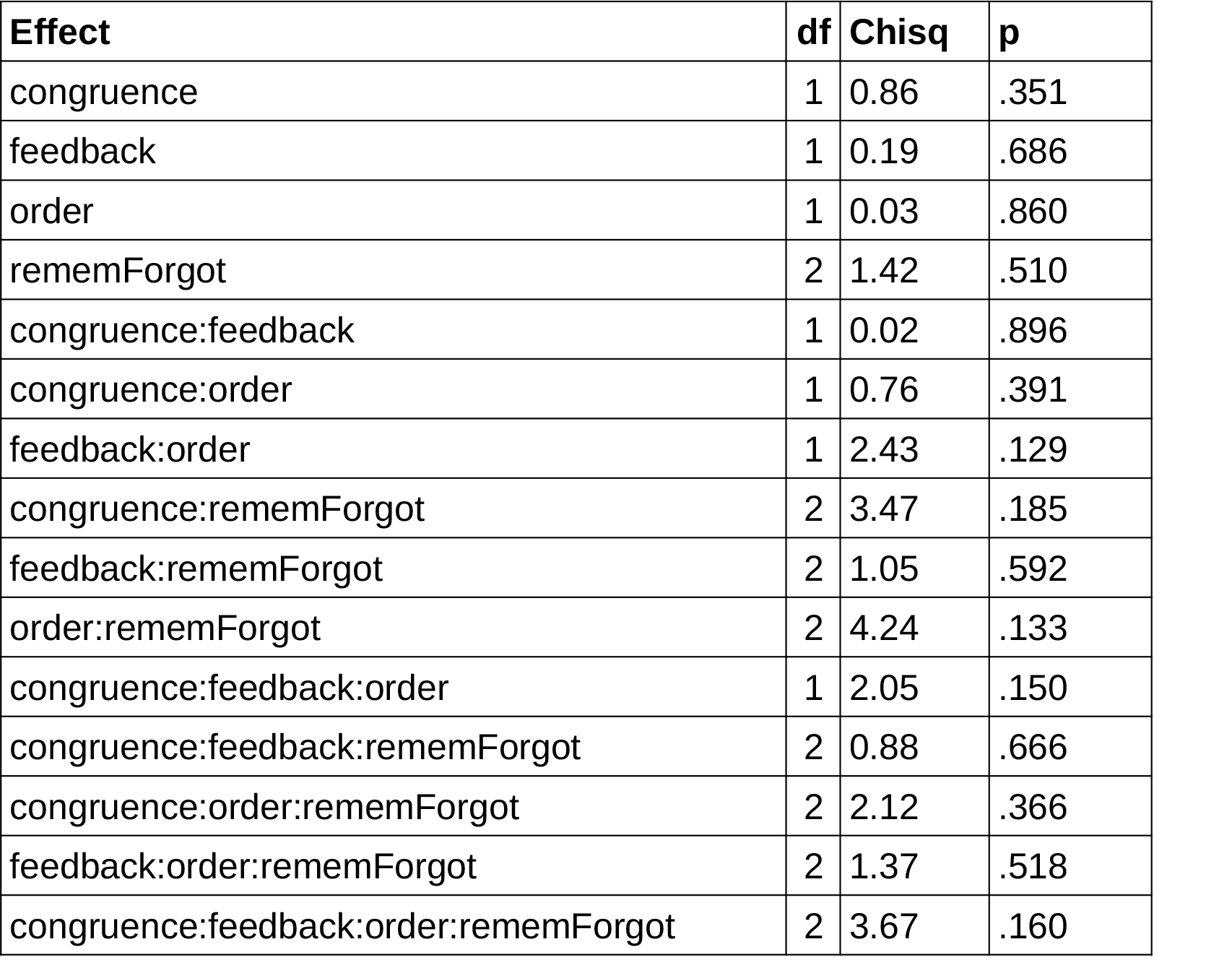
Mixed ANOVA statistical results for arousal ratings on day 2 based on performance on day 2 (remembered/forgotten). Mixed models were estimated with *congruence*, *feedback*, *feedback-order* and *rememForgot* as independent variables. Note that Factor “*rememForgot*” has 3 levels: Remembered (correct on both days), forgotten (correct only on day 1) or incorrect (already on day1 incorrect). A random intercept per subject was included. P-values were calculated with parametric bootstrapping. df = degrees of freedom; Chisq = chi square value; p = probability. “X:Y” denotes the interaction of X with Y.

| Effect | df | Chisq | p |
| --- | --- | --- | --- |
| congruence | 1 | 0.86 | .351 |
| feedback | 1 | 0.19 | .686 |
| order | 1 | 0.03 | .860 |
| rememForgot | 2 | 1.42 | .510 |
| congruence:feedback | 1 | 0.02 | .896 |
| congruence:order | 1 | 0.76 | .391 |
| feedback:order | 1 | 2.43 | .129 |
| congruence:rememForgot | 2 | 3.47 | .185 |
| feedback:rememForgot | 2 | 1.05 | .592 |
| order:rememForgot | 2 | 4.24 | .133 |
| congruence:feedback:order | 1 | 2.05 | .150 |
| congruence:feedback:rememForgot | 2 | 0.88 | .666 |
| congruence:order:rememForgot | 2 | 2.12 | .366 |
| feedback:order:rememForgot | 2 | 1.37 | .518 |
| congruence:feedback:order:rememForgot | 2 | 3.67 | .160 |

The arousal ratings on day 2 did not cause any many effects or interactions (all p’s >.13, see STab. 6).

## Confidence on day 2 based on remembered/forgotten

**Supplemental Fig. 7:**
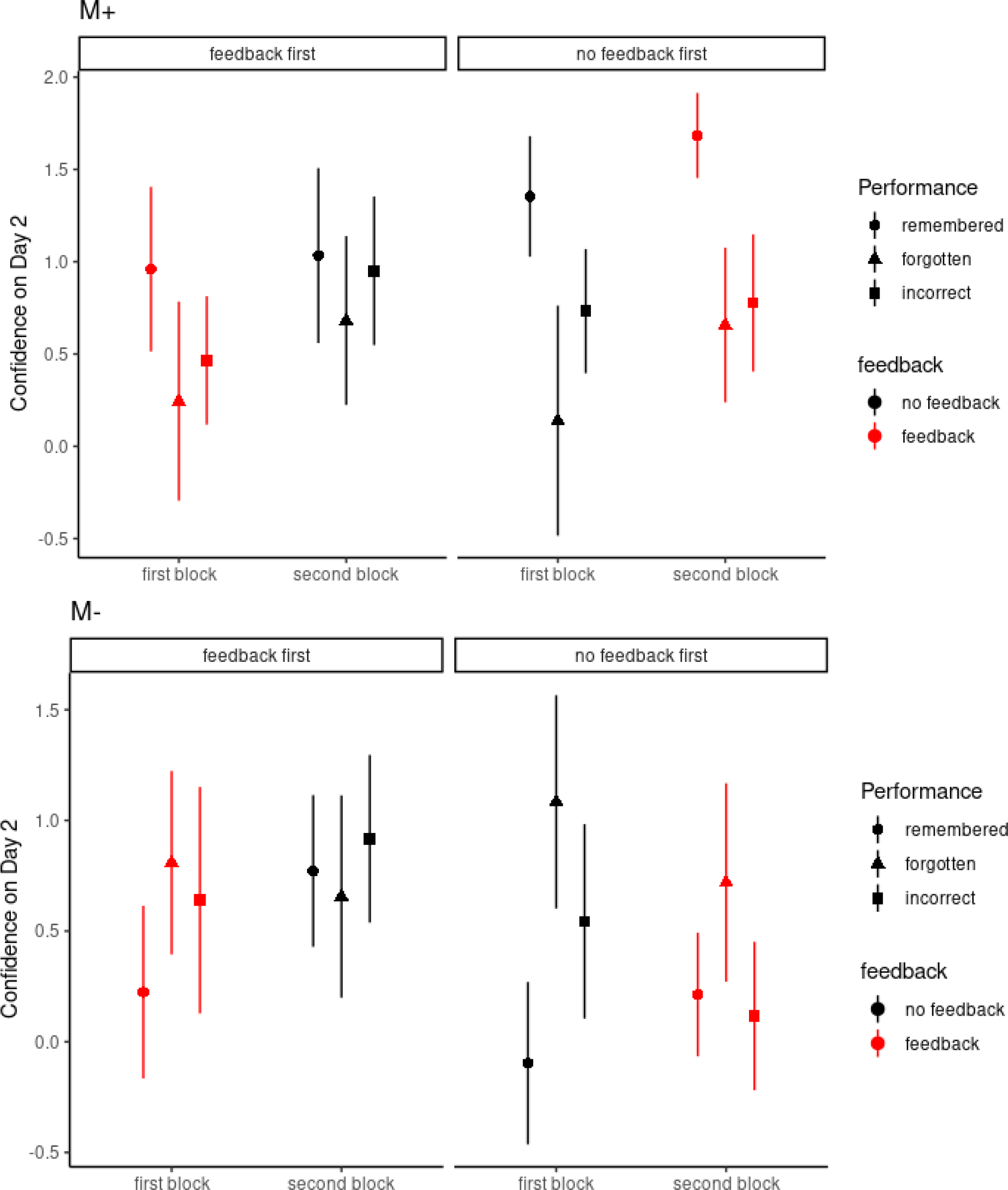
Condition-specific group means and between-subject SEM for Confidence ratings on day 2 sorted by performance on day 2 (remembered/forgotten/incorrect). Upper panel: M+ (dopamine-dependent trials). Lower panel: M- (dopamine independent trials). Left part of graphs: feedback first group; Right Part of graphs : no feedback first group.

**Supplemental Table 7:**
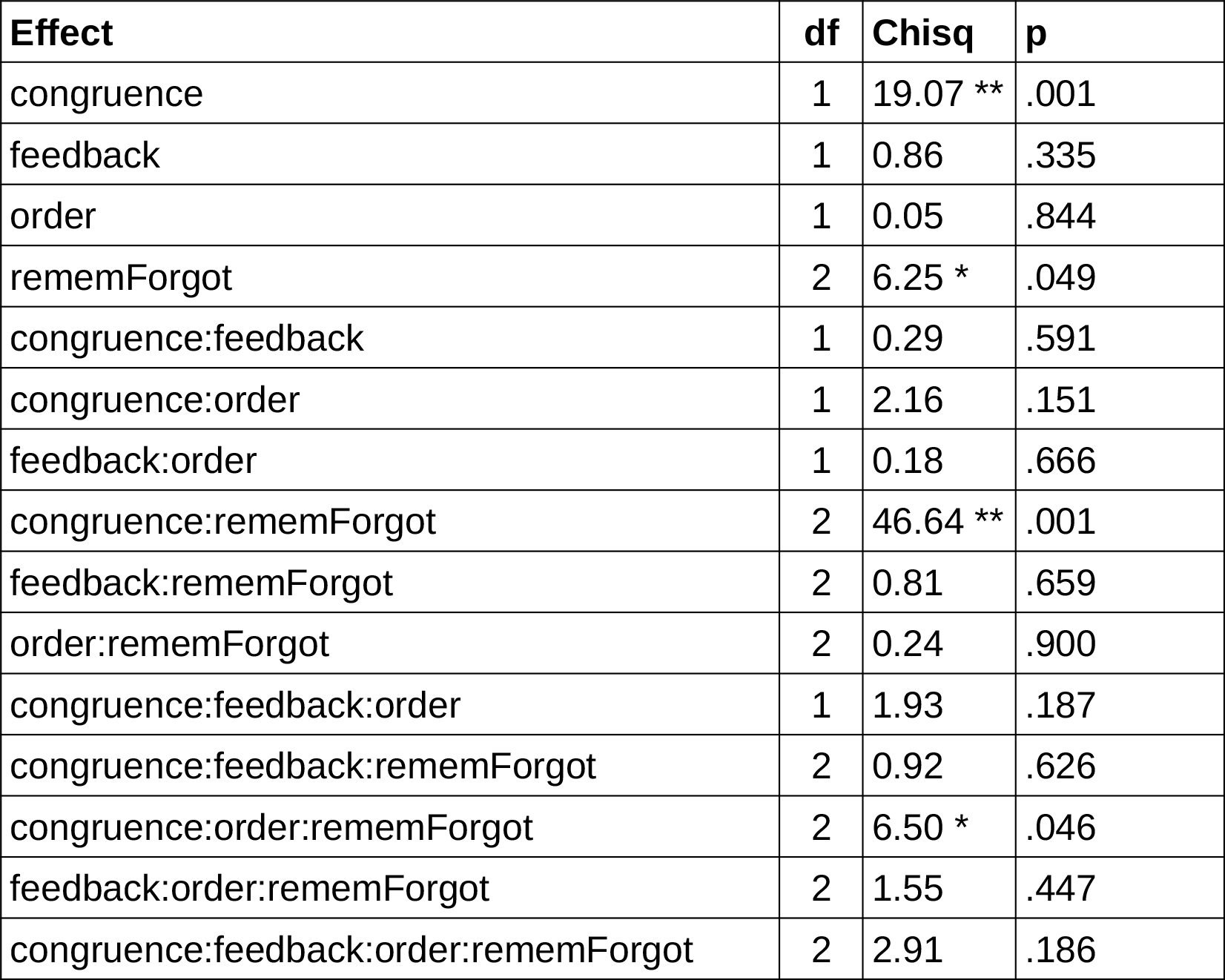
Mixed ANOVA statistical results for confidence ratings on day 2 based on performance on day 2 (remembered/forgotten). Mixed models were estimated with *congruence*, *feedback*, *feedback-order* and *rememForgot* as independent variables. Note that Factor “*rememForgot*” has 3 levels: Remembered (correct on both days), forgotten (correct only on day 1) or incorrect (already on day1 incorrect). A random intercept per subject was included. P-values were calculated with parametric bootstrapping. df = degrees of freedom; Chisq = chi square value; p = probability. “X:Y” denotes the interaction of X with Y.

| Effect | df | Chisq | p |
| --- | --- | --- | --- |
| congruence | 1 | 19.07 ** | .001 |
| feedback | 1 | 0.86 | .335 |
| order | 1 | 0.05 | .844 |
| rememForgot | 2 | 6.25 * | .049 |
| congruence:feedback | 1 | 0.29 | .591 |
| congruence:order | 1 | 2.16 | .151 |
| feedback:order | 1 | 0.18 | .666 |
| congruence:rememForgot | 2 | 46.64 ** | .001 |
| feedback:rememForgot | 2 | 0.81 | .659 |
| order:rememForgot | 2 | 0.24 | .900 |
| congruence:feedback:order | 1 | 1.93 | .187 |
| congruence:feedback:rememForgot | 2 | 0.92 | .626 |
| congruence:order:rememForgot | 2 | 6.50 * | .046 |
| feedback:order:rememForgot | 2 | 1.55 | .447 |
| congruence:feedback:order:rememForgot | 2 | 2.91 | .186 |

For confidence ratings we found significant main effects of C*ongruence* and *RememForgot*, their interaction and a triple interaction of *congruence*, *rememForgot* with *order*. This interaction was due to highest confidence rating in the no feedback first group for remembered M+ trials (relative to forgotten and incorrect trials) and lowest confidence ratings for the forgotten trials; for M-trials this pattern was reversed. In the feedback first group the overall pattern was similar yet attenuated.

